# DksA-Dependent Stringent Stress Response Drives Virulence and Gastrointestinal Persistence of *Klebsiella pneumoniae*

**DOI:** 10.64898/2026.03.18.712580

**Authors:** Md. Maidul Islam, Robert L. Beckman, Noah A. Nutter, Juan D. Valencia-Bacca, Giovanna E Hernandez, Renee M. Fleeman, Karen M Haas, M. Ammar Zafar

## Abstract

Successful gastrointestinal colonization (GI) by bacterial pathogens requires adaptation to nutrient competition and host-derived stresses in the gut, with adaptation via the bacterial stringent stress response playing a critical role. Epidemiological data suggest that the GI tract serves as a reservoir from where *K. pneumoniae* can spread and cause invasive disease or transmit to another host. DksA is a conserved stringent response transcriptional regulator that was identified in an *in vivo* transposon mutagenesis screen as an important *K. pneumoniae* gut determinant. However, its role in *K. pneumoniae* pathogenesis and gut colonization remains uncharacterized. Here, we demonstrate that DksA is required for survival against membrane-targeting antibiotics, consistent with a role in cell envelope stress tolerance. In addition, DksA positively regulates capsule biosynthesis gene expression and hypermucoviscosity and is essential for robust biofilm formation. Using a murine model, we show that DksA functions as a determinant of GI colonization independently of the resident gut microbiota. Furthermore, we demonstrate that DksA is important for environmental survival and transmission by regulating RpoS, thereby providing a mechanistic link between the stringent stress response, environmental survival, and subsequent transmission. Together, these findings establish DksA as a central integrator of the stringent response, coordinating membrane stress resistance, virulence traits, and gastrointestinal colonization in *K. pneumoniae*.

**Importance:** *K. pneumoniae,* a pathobiont, is responsible for multidrug-resistant infections and poses a major threat in hospital settings as well as community-acquired invasive infections. The bacterium tightly coordinates its virulence-associated traits to adapt to diverse environmental conditions and survive; however, the regulatory mechanisms remain poorly understood. In this study, we demonstrated that the conserved stringent response regulator DksA contributes to bacterial membrane stability, thereby affecting antibiotic resistance, inherent virulence, and persistence traits of *K. pneumoniae*. Additionally, DksA was identified as required for gut colonization, environmental survival through dysregulation of RpoS, and transmission to a naïve host. These results enhance our overall understanding of *the K. pneumoniae* stringent response and will provide new avenues for controlling *K. pneumoniae* infections.

## Introduction

The human gut is a competitive environment in which both the host and the microbiota vie for nutrients. This competition for resources results in inhibition of colonization by enteric pathogens, a phenomenon known as colonization resistance (1). Because of the intense competition, gut commensal organisms dedicate significant resources to nutrient acquisition. Gut bacteria undergo periods of famine and feast, requiring rapid responses that are generally mediated by transcriptional regulators and small molecules (alarmones) that allow them to respond to drastic nutritional changes as a consequence of the host diet (2).

*Klebsiella pneumoniae* is a gram-negative pathobiont that frequently colonizes mammalian mucosal surfaces, including the gastrointestinal (GI) tract (3). From this mucosal site *K. pneumoniae* can spread from the GI tract and cause invasive disease in the same host such as urinary tract infections (UTI), bloodstream infections, liver abscesses, and pneumonia (4, 5), as well as transmit to a new host. The worldwide dissemination of multidrug-resistant (MDR) and hypervirulent pathotypes of *K. pneumoniae* poses a major challenges for both treatment and control (6). Recent studies have highlighted how *K. pneumoniae* elicits a subclinical inflammatory response, utilizes alternative nutrient sources and direct inhibition of the gut microbiota as molecular pathways it employs to overcome colonization resistance CR (7–10). Similar to members of the gut microbiota, *K. pneumoniae* likely encounters stresses, including but not limited to antimicrobial peptides, changes in osmotic and pH level, and nutrient competition.

The mechanism of bacterial adaptation in response to nutrient deprivation and other stresses is known as stringent response (11). Bacterial stress regulating systems are complex and involve a large number of cellular processes to fine tune, allowing for cell survival and proliferation (12).

One such response is the bacterial stringent response that has been implicated in regulation of virulence genes, antibiotic tolerance (13, 14) and survival of certain commensals in the gut (2, 15). During stringent response, gene expression is mainly controlled by guanosine penta- or tetra-phosphate (collectively referred to as (p)ppGpp), synthesized by RelA and SpoT (16, 17). (p)ppGpp working in conjunction with the small conserved protein DksA, causes conformational changes to RNA polymerase (RNAP) resulting in profound changes in gene expression (18, 19). The modulation of gene expression by (p)ppGpp and DksA can be positive or negative depending on the properties of the promoters (20). In addition, the interaction between (p)ppGpp and DksA for transcription can be synergistic or antagonistic (21, 22).

A recent *in vivo* transposon mutagenesis screen to identify *K. pneumoniae* gut determinants included the transcriptional regulator DksA (7), which is highly conserved in Gammaproteobacteria, with transcriptomic analyses revealing that it regulates approximately 5-20% genes of the genome (23, 24). DksA has been implicated in diverse bacterial processes, including bacterial division, virulence regulation, quorum sensing, intestinal colonization and antibiotic tolerance is diverse pathogens including *Salmonella enterica* (25, 26), *Pseudomonas aeruginosa* (27), *Vibrio cholerae* (28), *Shigella flexneri* (29), *Escherichia coli* (30, 31),

*Acinetobacter baumanni* (32, 33) and *Yersinia enterocolitica* (34, 35). Collectively, these findings establish DksA as a global regulator that coordinates stringent response, bacterial physiology, and pathogenicity across diverse species. However, its role in *K. pneumoniae* remains to be elucidated.

Herein, to provide an understanding of the molecular pathways that DksA regulates in *K. pneumoniae*, we investigated its role in coordinating antibiotic tolerance, key virulence traits, and intestinal colonization. We demonstrate that DksA is important for survival against membrane-targeting antibiotics, positively regulates hypermucoviscosity and capsule production, and critical for biofilm development. We further show that DksA is required for gut colonization independent of the microbiota and contributes to survival outside the host and subsequent transmission. These findings suggest that DksA functions in a master regulatory role in virulence and persistence pathways, thereby promoting both pathogenesis and gut colonization of *K. pneumoniae*.

## Results

DksA is required for optimal growth of *K. pneumoniae* in minimal medium and survival against membrane targeting antibiotics.

As DksA was initially identified as a *K. pneumoniae* gut determinant in a transposon mutagenesis screen (7), and is involved in stringent stress response including growth, we examined its contribution to *K. pneumonia* growth in nutrient-rich and limited (defined) media. We compared the growth of KPPR1S (a derivative of ATCC 43816; wild type [WT]) (**Table. 1**) to its isogenic mutant *dksA::cam* (*dksA^-^*) and the chromosomally complemented (reconstituted at the native site) strain *dksA^+^*. We first assessed the growth using nutrient-rich media (lysogeny broth [LB]) and M9 minimal media (MM) supplemented with glucose which served as defined media. While all the strains grew similarly in LB (**Fig. S1A**), the *dksA^-^* strain grew poorly in M9 MM (**Fig. S1B**). However, the addition of 0.2% Casamino Acids (CAA) led to full growth recovery of the mutant (**Fig. S1C**). Our growth findings with *K. pneumoniae* are comparable to other related species that highlight the importance of *dksA* in nutrient limited conditions (25, 27, 36). Next, a transcriptional *gfp* fusion with the *dksA* promoter was used to determine the expression of *dksA* during *K. pneumoniae* growth in LB, where we observed a growth phase dependent increase in GFP levels (**Fig. S1D**). Similar to *E. coli*(37), GFP levels were elevated in the *dksA^-^* strain (**Fig. S1D**) indicating that negative autoregulation of DksA also occurs in *K. pneumoniae* at the transcriptional level.

**Table 1.** Strains and plasmids used in the study.

| Strain | Description | Antibiotic resistance phenotype | Reference or source |
| --- | --- | --- | --- |
| AZ16 | *E. coli* S17-1 λpir<br>Donor strain for conjugation<br>*RP4::2-Tc::Mu::Km Tn7* | Tp<sup>r</sup>; Sm<sup>r</sup> | (96) |
| AZ55 | KPPR1S (ATCC43816), Str<sup>r</sup><br>derivative of KPPR1 | Rif<sup>r</sup>, Str<sup>r</sup> | (91) |
| AZ63 | AZ55 with pKD46 plasmid for<br>lambda red recombination | Rif<sup>r</sup>, Str<sup>r</sup>, Spec<sup>r</sup> | (40) |
| AZ94 | Derivative of KPPR1 with<br>apramycin resistance cassette at<br>*att::Tn7* site | Rif<sup>r</sup>, Apra<sup>r</sup> | (97) |
| AZ359 | Arrayed library MKP103 with<br>transposon element in *dksA::cam*,<br>(strain: tnkp1_lr150117p10q173) | Cam<sup>r</sup> | (84) |
| AZ360 | *dksA::cam* from AZ359 into AZ55 | Rif<sup>r</sup>, Str<sup>r</sup>, Cam<sup>r</sup> | This study |
| AZ387 | $\Delta dksA$, clean deletion created from<br>AZ360 using pFlp3; plasmid cured | Rif<sup>r</sup>, Str<sup>r</sup> | This study |
| AZ388 | *dksA*<sup>+</sup> (chromosomally<br>complemented) derivative of<br>AZ387 | Rif<sup>r</sup>, Str<sup>r</sup> | This study |
| AZ434 | AZ55 with gfp at *att::Tn7* site | Str<sup>r</sup>, Rif<sup>r</sup>, Amp<sup>r</sup>,<br>Kan<sup>r</sup> | This study |
| AZ436 | AZ360 with gfp at *att::Tn7* site | Str<sup>r</sup>, Rif<sup>r</sup>, Amp<sup>r</sup>,<br>Kan<sup>r</sup>, Cam<sup>r</sup> | This study |
| AZ438 | AZ388 with gfp at *att::Tn7* site | Str<sup>r</sup>, Rif<sup>r</sup>, Amp<sup>r</sup>,<br>Kan<sup>r</sup> | This study |
| **Plasmids** |  |  |  |
| pKD46 | $\lambda$ red recombinase genes<br>downstream of *araBAD* promoter;<br>temperature sensitive (32°C) | Spec<sup>r</sup> | (98) |
| pCre2 | Flippase's for removing *cam*<br>cassettes | Amp<sup>r</sup> | (99) |
| pKAS46 | Vector for allelic exchange,<br>contains *rpsL* for streptomycin<br>counter-selection | Kan<sup>r</sup>, Str<sup>s</sup> | (86) |
| pPROBE | gfp transcriptional reporter vector | Kan<sup>r</sup> | (88) |
| pPROBE_*rmpA* | *rmpA* promoter region of KPPR1S<br>in pPROBE | Kan<sup>r</sup> | (72) |
| pPROBE_*galF* | *galF* promoter of KPPR1S in<br>pPROBE | Kan<sup>r</sup> | (44) |
| pPROBE_*wzi* | *wzi* promoter of KPPR1S in pPROBE | Kan<sup>r</sup> | (44) |
| pPROBE_*manC* | *cpsB* promoter of KPPR1S in pPROBE | Kan<sup>r</sup> | (44) |
| pPROBE_*mrkA* | *mrkA* promoter of KPPR1S in pPROBE | Kan<sup>r</sup> | This study |
| pPROBE_*dksA* | *dksA* promoter of KPPR1S in pPROBE | Kan<sup>r</sup> | This study |
| pUC18R6K-mini-Tn7T-Km | R6K origin of replication which needs *pir* gene for replication | Kan<sup>r</sup>, Amp<sup>r</sup> | (90) |
| pTNS2 | Tn7 transposase expression | Amp<sup>r</sup> | (90) |
| pJH026 | pPROBE with constitutive *pem7* promoter | Kan<sup>r</sup> | (89) |

Rif<sup>r</sup>, rifampicin resistant, Str<sup>r</sup>, streptomycin resistant, Apra<sup>r</sup>, apramycin resistant, Cam<sup>r</sup>, chloramphenicol resistant, Spec<sup>r</sup>, spectinomycin resistant, Amp<sup>r</sup> ampicillin resistant, Kan<sup>r</sup>, kanamycin resistant, Str<sup>s</sup>, streptomycin sensitive.

As DksA mediated stringent stress response is implicated in antibiotic resistance (34, 38), we investigated whether *K. pneumoniae dksA* affects antibiotic resistance. We tested survival following exposure to different classes of antibiotics, including cationic antimicrobial peptide, aminoglycoside, cephalosporin, quinolone and penicillin. Antimicrobial peptides such as polymyxin B disrupt the lipopolysaccharide present in the outer-membrane leading to membrane depolarization and eventual lysis of the inner membrane (39). Compared to the WT strain, which is sensitive to polymyxin B (40), the *dksA^-^* strain exhibited enhanced resistance to increasing concentrations of polymyxin B (**Fig. 1A**). The chromosomally complemented *dksA^+^* strain behaved similarly to the WT strain, suggesting that the observed higher resistance is specific to the deletion of *dksA*. In addition, loss of *dksA* resulted in increased survival against the aminoglycoside antibiotics tobramycin and kanamycin (**Fig. 1B, C**). However, we observed no distinction in resistance profile among the WT, mutant, and the complement strain when treated with cefixime, nalidixic acid and oxacillin (**Fig. 1D**, and **Fig. S2**).

**Figure 1:**
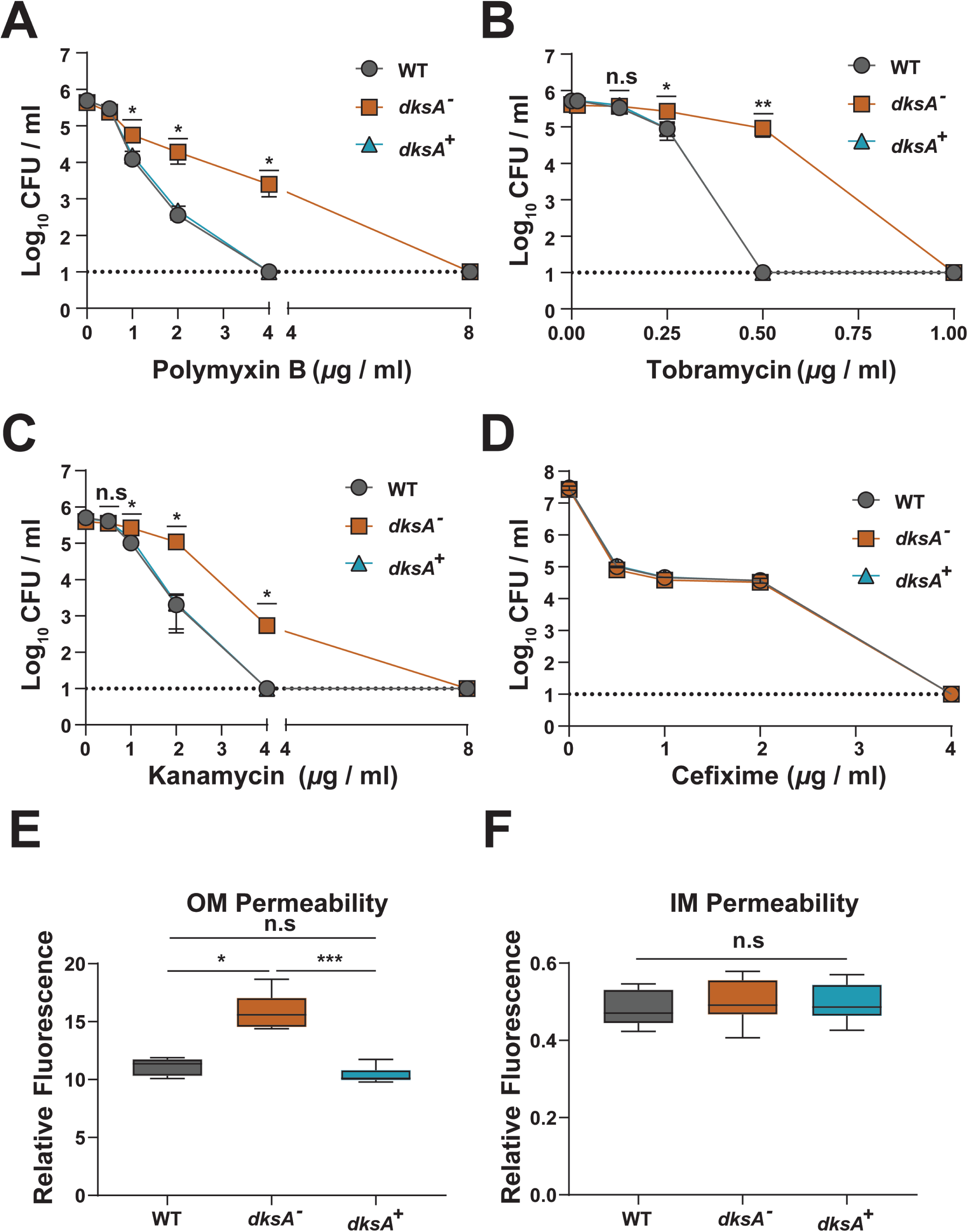
DksA is required for *K. pneumoniae* survival against membrane targeting antibiotics. **(A-D)** WT, *dksA*^-^, and *dksA^+^* strains were grown until OD_600_=∼1.0 in cation-adjusted MHB and diluted to ∼10^6^ CFU/mL in fresh caMHB. Diluted samples were mixed with appropriate concentration of polymyxin B **(A)**, Tobramycin **(B)**, Kanamycin **(C)**, Cefixime **(D)** and incubated for 30 minutes (for cefixime the incubation time was 3 hours) at 37°C. Shown is the mean and SEM of three independent assays (in triplicate). **(E-F)** Relative fluorescence intensity for outer membrane (OM) and inner membrane (IM) permeability assays, detected by fluorescent dye N-phenyl-1-naphthylamine (NPN) and propidium iodine (PI) uptake, respectively. Statistical differences were calculated using Kruskal-Wallis tests with Dunn’s test of multiple comparisons within each concentration group. There was no significant difference between the strains at different concentrations of cefixime. *\**, *P <* 0.05; \**\**, *P <* 0.01; \*\**\**, *P <* 0.001; n.s., not significant.

The modification in antibiotic resistance profile of *K. pneumoniae* to polymyxin B led us to investigate the role of *dksA* in structural and functional integrity of the cell membrane. To determine the integrity of the cell membrane, the permeability of the outer membrane (OM) and inner membrane (IM) were measured by N-phenyl-1-naphthylamine (NPN) and propidium iodide (PI) uptake assay respectively. NPN is a hydrophobic fluorescent probe which binds hydrophobic structures (phospholipids) in the OM resulting in an increase in fluorescence. We observed an increase in fluorescence with the *dksA^-^* strain compared to WT (**Fig. 1E**). Permeability of the IM was measured by a PI uptake assay. PI binding to nucleic acid triggers fluorescence, but it can only gain access to the cytoplasm when the outer and inner membrane are damaged. In contrast to our data with NPN, which indicated a defect in OM we did not observe a difference in PI mediated fluorescence among the tested strains, suggestive of an intact IM (**Fig. 1F**). Together, these findings illustrate that, while *K. pneumoniae* DksA, like other Gammaproteobacteria, contributes to growth, its role in antibiotic resistance is comparatively more disparate.

### DksA positively regulates capsule gene expression and HMV production in *K. pneumoniae*

Previous transposon mutagenesis screens implicated DksA in capsular polysaccharide (CPS) production (41, 42), a key *K. pneumoniae* virulence determinant. Additionally, *K. pneumoniae* CPS sensitizes strains to polymyxins(43), suggesting that, in addition to outer membrane integrity, DksA-dependent changes in CPS levels might also contribute to the observed polymyxin resistance (**Fig. 1A**). Thus, to assess the contribution of DksA in regulating CPS we utilized three transcriptional fusion plasmids containing promoter regions of *galF*, *wzi*, or *manC* genes that encode enzymes involved in CPS production and export (**Fig. 2A**) (44).

**Figure 2:**
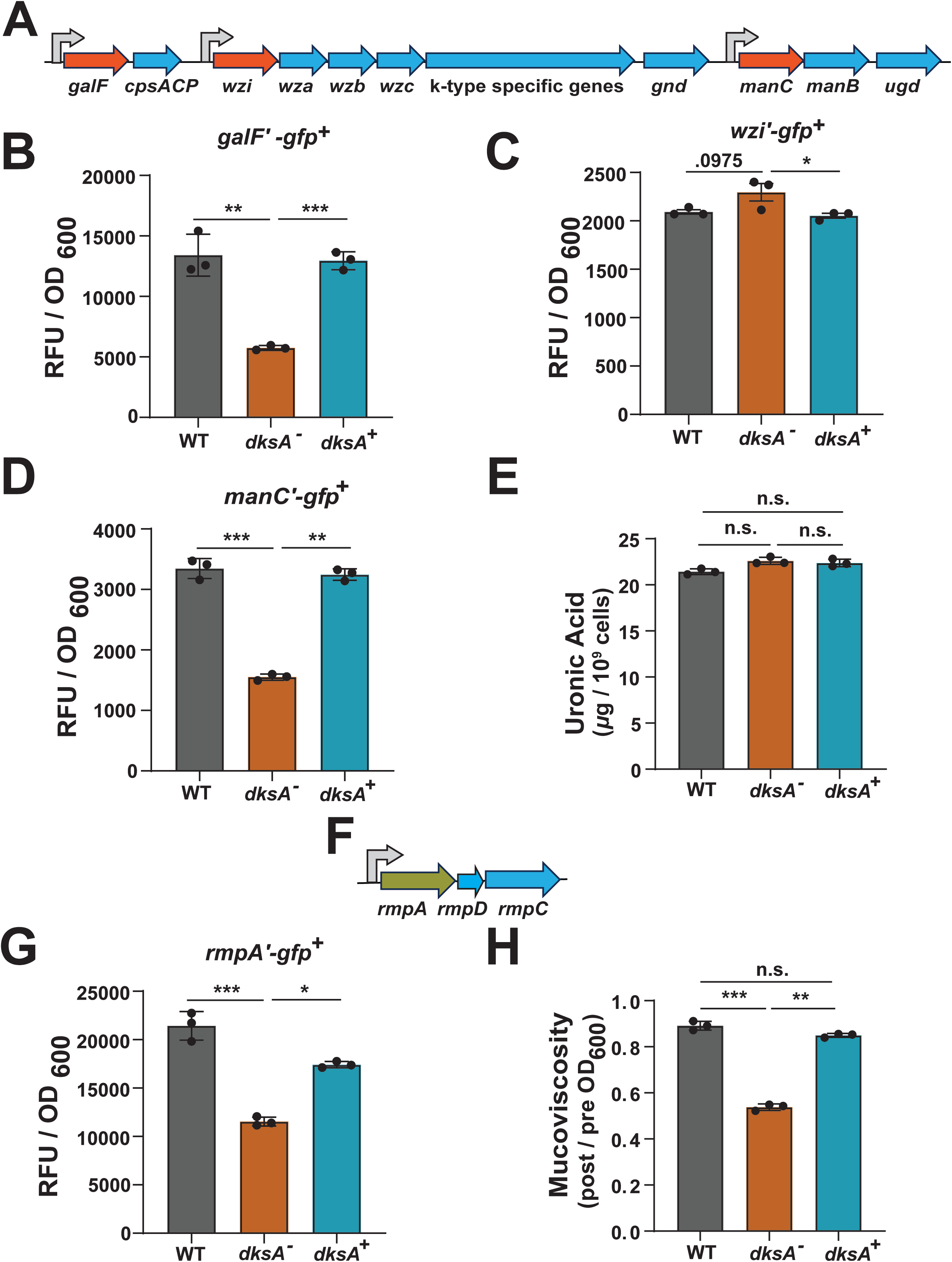
DksA positively influences hypermucoviscocity but not capsule production. **(A)** Genetic organization of the capsule locus of *K. pneumoniae* strain KPPR1. Promoter elements of the genes highlighted in orange were fused to *gfp*. **(B-D)** *gfp* expression assay using WT, *dksA*^-^, and *dksA^+^* strains carrying plasmids with promoter regions from capsule locus fused to *gfp,* (**A)** *galF* promoter (pPROBE-*galF*′*-gfp^+^*), **(B)** *wzi* promoter (*wzi*′*-gfp^+^*), **(C)** *manC* promoter (*manC*′*-gfp^+^*). Overnight samples were adjusted to OD_600_ = 0.2 in fresh LB and incubated in rotor for 6 hours at 37°C and the relative fluorescent units (RFU) were determined and normalized to OD_600_. **(E)** Uronic acid quantification was used to measure capsule production in WT, *dksA*^-^, and *dksA^+^* from samples grown to stationary-phase. **(F)** Genetic organization of the *rmp* locus of *K. pneumoniae* strain KPPR1. Promoter element of the gene highlighted in green was fused to *gfp*. **(G)** *gfp* expression assay using WT, *dksA*^-^, and *dksA^+^*strains carrying plasmid with *rmp* promoter transcriptional *gfp* fusion(*rmpA*′*-gfp^+^*). **(H)** Comparison of the hypermucoviscosity (HMV) between WT, *dksA*^-^, and *dksA^+^* using the sedimentation assay. All the data are represented from 3 independent assays (n≥3). Kruskal-Wallis tests with Dunn’s test of multiple comparisons were performed on the pooled samples across strains to determine statistical significance. *\**, *P <* 0.05; \**\**, *P <* 0.01; \*\**\**, *P <* 0.001; n.s., not significant.

While we observe significantly reduced *gfp* expression from the *galF* and *manC* promoters in *dksA^-^* strain, with complementation of *dksA* (*dksA^+^*) restoring GFP levels (**Fig. 2B, D**), there was slight increase in the GFP levels from the *wzi* promoter in *dksA^-^* strain compared to WT strain (**Fig. 2C**). While we observed differences at the transcriptional level at two out of three promoters, we did not observe corresponding differences in uronic acid (UA) content, a key component of *K. pneumoniae* CPS, between WT, *dksA^-^*and *dksA^+^* (**Fig. 2 E**). These results highlight that the gene expression from the *wzi* promoter, involved in CPS production and export, likely compensates for reduced expression from the other two CPS promoters, resulting in similar UA levels between the WT and the *dksA*^-^ strain.

Next, we focused on hypermucoviscous (HMV) phenotype, which is distinct from CPS production, with the *rmp* locus (**Fig. 2F**) driving the HMV phenotype (45). Therefore, we ascertained the role of DksA on the expression of *rmpA* promoter. Compared to the WT and the *dksA*^+^ strain, we observed a 2-fold reduction in GFP levels from the *rmpA* promoter in the *dksA^-^*background (**Fig. 2G**). This reduction in *gfp* expression from the *rmp* promoter was concomitant with a reduced HMV levels in the *dksA^-^*strain (**Fig. 2H**). All together these results indicate that DksA of KPPR1S is an important determinant for the HMV phenotype but does not alter the CPS UA content.

### DksA is required for robust biofilm formation by *K. pneumoniae*

As our data demonstrated DksA regulating mucoviscosity, we theorized that DksA might also modulate the expression of *K. pneumoniae* factors required for biofilm formation, another critical *K. pneumoniae* virulence determinant. Consequently, we performed biofilm formation assay using a microtiter dish colorimetric assay. Compared to the WT and *dksA*^+^, the *dksA^-^* strain formed 7-fold less biofilm when grown in TSB with 0.5% glucose, (**Fig. S3A**) thus highlighting the importance of DksA in biofilm formation. While our microtiter plate biofilm assay allowed us to assess the contribution of DksA to *K. pneumoniae* biofilm biomass, it did not provide insight into DksA impact on biofilm structural matrix. For this purpose, we used strains with *gfp* constitutively expressed off the chromosome (*attTn7::gfp*) which were grown in two different conditions: static condition and microfluidic condition (flow rate of 65 µL h^−1^ for 24-h to resembles environmental shear flow) as described by Beckman et al. (46). Biofilm matrix was evaluated using confocal z-stack imaging, using GFP fluorescence to visualize the cells, Nile red to detect the presence of lipids (red) and calcofluor white (blue) to visualize the polysaccharide matrix. These parameters allowed us to visualize the 3D spatial distribution of cellular populations, and the lipid and polysaccharide content distribution within the matrix. In both static and microfluidic conditions, the WT strain formed robust (∼30 µm) biofilm whereas the *dksA^-^* strain had reduced (10 µm) biofilm height as shown in confocal z-stack imaging, with the *dksA*^+^ strain restoring the WT phenotype (**Fig. 3** and **Fig. S4**). Furthermore, we observed a thick polysaccharides matrix with WT cells present above and below the layer, whereas *dksA^-^* strain had a thin layer of polysaccharide matrix with less cells present outside of the matrix.

**Figure 3:**
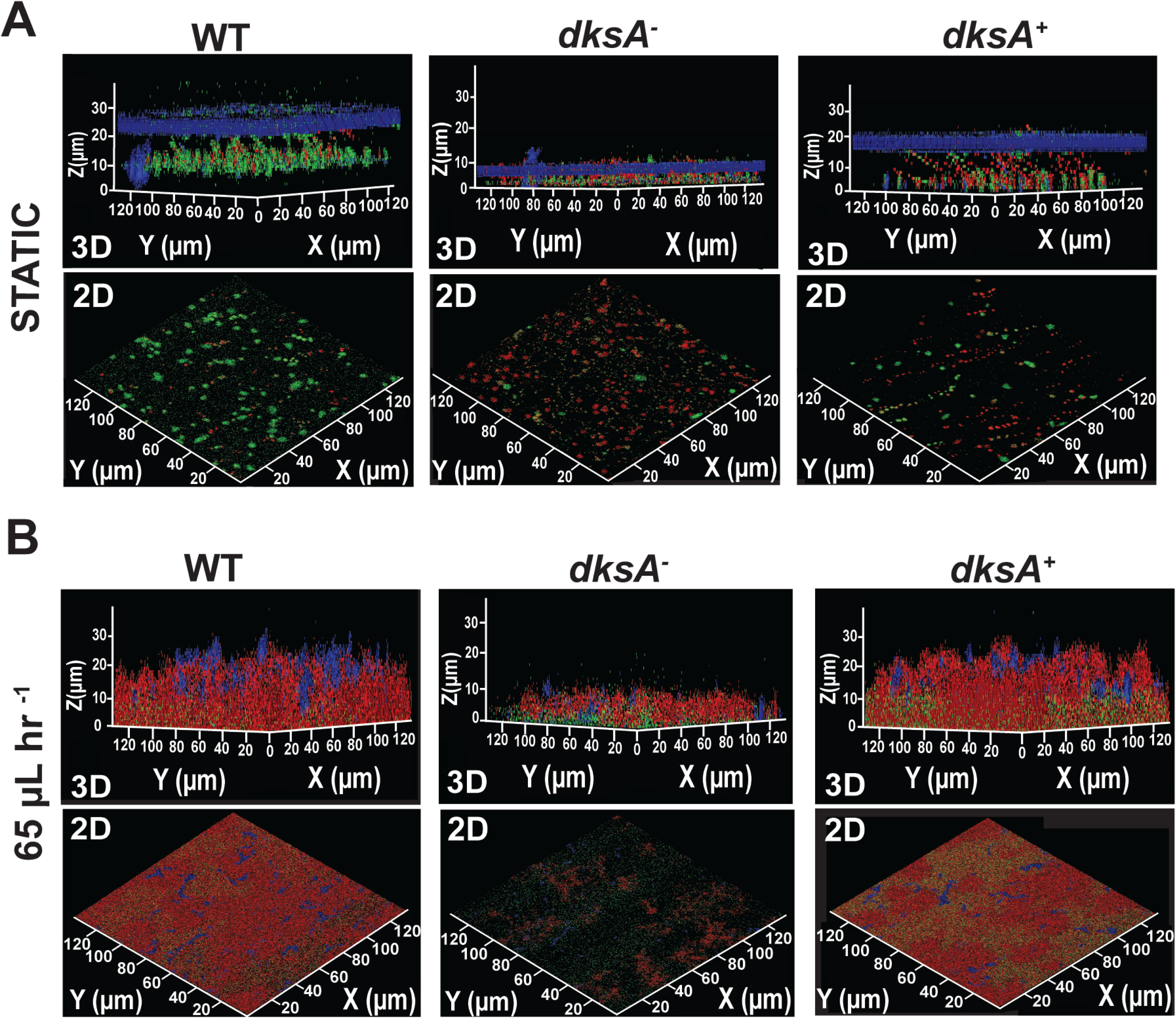
Biofilm matrix components are positively regulated by DksA resulting in robust biofilm formation. Shown are the 2D and 3D rendering of confocal z-stack imaging of chromosomally encoding GFP *K. pneumoniae* WT, *dksA*^-^, and *dksA^+^* strains, Nile red to visualize the presence of lipids (red) and calcofluor white (blue) to visualize the polysaccharide matrix, respectively. Each strain shown is either grown under static conditions in biofilm media (Tryptic soy broth with 0.5% glucose) at 37°C **(A)** or with 65 μl h^-1^ shear flow **(B)**. For each strain and the two conditions the z-stack images are shown to visualize the height of the biofilm and images from the top of the biofilm are shown to visualize the overall cellular density. Biofilms were grown and imaged in triplicates with a representative image shown.

Collectively, our biofilm assays build on each other and provide insight in to *K. pneumoniae* biofilm architecture and identify DksA as likely regulating multiple biofilm matrix parameters that are required for robust biofilm formation.

Subsequently, we sought to identify DksA regulated molecular pathways that control *K. pneumoniae* biofilm formation. As type 3 fimbriae (**Fig. S3B**) promotes biofilm formation in *K. pneumoniae* (9, 47), we examined whether DksA regulated the expression of type 3 fimbriae genes *mrkA* and *mrkB* through quantitative reverse transcription-PCR (qRT-PCR). Compared to the WT, the transcript levels of both *mrkA* and *mrkB* genes were significantly reduced in the *dksA^-^* strain (**Fig. S3B**). Consistent with the qRT-PCR data, our *mrkA’*-*gfp*^+^ transcriptional fusion fluorescence results also showed reduced *gfp* expression in the *dksA^-^* strain (**Fig. S3C**). In addition to type 3 fimbriae, quorum sensing (QS) has been linked to biofilm formation, with *K. pneumoniae* primarily using autoinducer-2 quorum sensing (AI-2 QS) system which is synthesized by the *luxS* gene (48–50). *K. pneumoniae* regulates its AI-2 QS signaling through *lsrACDBFG* and *lsrRK* operons (51). Our qRT-PCR results showed that *lsrR* and *lsrA* expression was significantly decreased in the *dksA^-^* strain compared to WT (**Fig. S3D**), suggesting that DksA potentially regulates multiple molecular pathways that are critical for biofilm formation.

### DksA is a gut colonization determinant independent of the resident gut microbiome

As *dksA* was identified in a mutagenesis screen as a *K. pneumoniae* gut colonization determinant (7) and observed to regulate several prominent virulence traits (**Fig. 2-3**) we decided to determine its role in gut colonization using our mouse model with an intact microbiota (52). Mice were orally inoculated with WT, *dksA^-^* and *dksA^+^* strains and then fecal pellets were collected daily to enumerate bacterial shedding, a non-invasive metric of GI colonization. During the experimental period (7 days), mice inoculated with *dksA^-^* strain shed poorly compared to WT and *dksA^+^* strains (**Fig. 4A**). Bacterial burdens in the oropharnynx and the lower intestine showed a similar trend (**Fig. S5A**). Subsequently, we performed a competition study using 1:1 mixture of WT and *dksA^-^* strain to investigate whether the WT would rescue the gut colonization defect of *dksA^-^* strain. However, the WT strain outcompeted the *dksA^-^* strain in the murine gut (**Fig. 4B**, **C**), suggesting that the WT strain is unable to rescue the gut colonization defect of the *dksA^-^*strain. In addition, we also performed a competition experiment between the *dksA^+^* and the WT strain and observed no difference in shedding and colonization between the strains (**Fig. S5B**, **C**), indicating that the defect in gut colonization was specific to loss of function of *dksA*.

**Figure 4:**
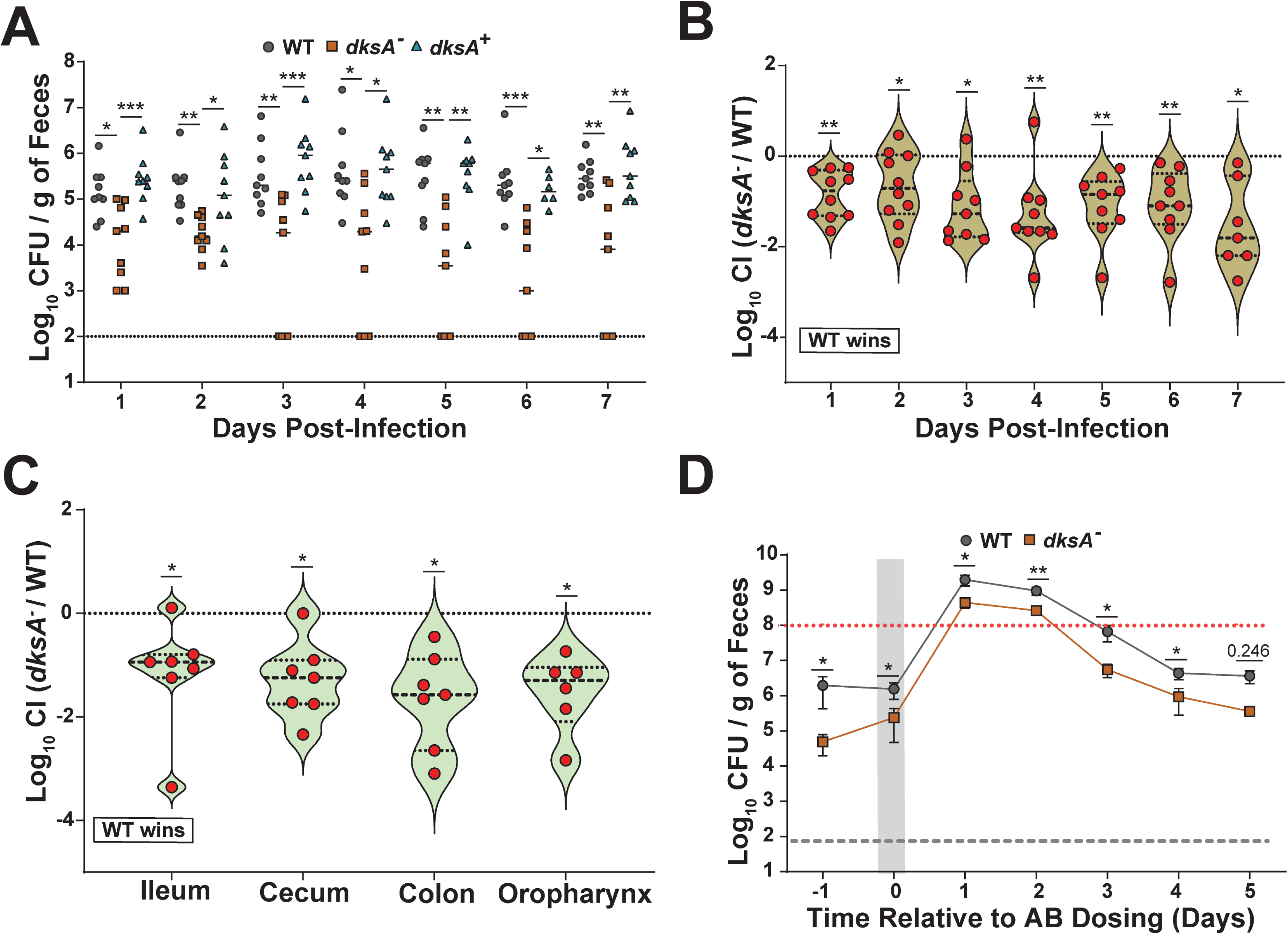
DksA is a gut colonization determinant during microbiota facilitated colonization resistance and during antibiotic dysbiosis. **(A)** Fecal shedding of *K. pneumoniae* infected mice. Mice were orally infected with 10^6^ CFU with either WT, *dksA*^-^, or *dksA^+^*strains (*n* = 9 for each group). Fecal pellets were collected daily from infected mice and *K*. *pneumoniae* was enumerated in the feces. Each symbol indicates a single mouse on a given day, the bars indicate the median shedding, and the dotted line indicates the limit of detection; significance is shown between the WT and *dksA^-^*, and *dksA^+^* and *dksA^-^*. Kruskal-Wallis test followed by Dunn’s test of multiple comparisons was performed at each time point for analysis. WT and *dksA*^+^ were not significantly different. (B) *K. pneumoniae* shedding from fecal sample from *in-vivo* competition index (CI) studies. Mice were infected orally with a 1:1 mixture of the WT and *dksA*^-^, with feces collected for 7 days (*n* =10). The CI was determined as described in Materials and Methods. Each point represents the log_10_ competitive index value from an individual mouse on the indicated day. Values above the dashed line indicate the mutant outcompeted the WT, whereas values below the dashed line indicate WT outcompeted the mutant. Bars indicate median value. The dashed line indicates a CI of 1 or a 1:1 ratio of WT to mutant. **(C)** Colonization density of the ileum, cecum, colon, and oropharynx represented as log_10_ CI values from an individual mouse at the end of the study. For both B and C, statistical significances were calculated using Wilcoxon signed-rank test with a theoretical median of 0. **(D)** Fecal shedding from *K. pneumoniae* from mice treated with antibiotic (AB). Mice were infected orally and 2 days postinoculation, they were given streptomycin (5 mg/200 µl) via oral gavage. The gray area indicates the time point for antibiotic treatment. Shown are the means and standard error of the means for both WT and *dksA*^-^ infected mice (*n*= 5 for each strain), the grey dotted line indicates the limit of detection, and the red dotted line indicates the super shedder threshold (≥10^8^ CFU [SS Threshold]). Statistical differences were calculated using a Mann-Whitney *U* test at each time point. *\**, *P <* 0.05; \**\**, *P <* 0.01; \**\*\**, *P <* 0.001; n.s., not significant.

To provide insight into whether the colonization defect of the *dksA^-^*strain was due to the presence of the native gut microbiota (colonization resistance) or because of an intrinsic defect, we treated *K. pneumoniae* inoculated mice with a single dose of antibiotic, resulting in transient depletion of the gut microbiome (52). Mice mono-infected with either the WT or *dksA^-^* were gavaged with a single dose of streptomycin 2 days postinfection to induce the supershedder phenotype (>10^8^ CFU/g of feces). After disruption of native host-microbiota both strains became supershedder, although the WT strain continued to shed significantly higher compared to *dksA^-^*strain post-treatment, and as the microbiome reset itself (**Fig. 4D**). Together, these results indicate that *K. pneumoniae dksA* is an important determinant of gut colonization both in the presence and absence of gut microbiota.

### DksA-dependent regulation of RpoS links environmental survival to gut colonization

The persistence of bacteria on medical equipment serves as reservoirs for infections of nosocomial pathogens in a hospital setting with *K. pneumoniae* ability to form biofilms enhancing its ability to survive in the environment (53, 54). DksA of *A baumanni*, another nosocomial pathogen, contributes to survival in harsh environments including desiccation tolerance (32). As *dksA^-^* strain had a defect in gut colonization and biofilm formation, we decided to test the environmental survivability of our strains. We used a previously described (40) solid surface survival protocol using nitrocellulose membranes on agarose pads and determined viability of the WT, *dksA^-^*, and *dksA* ^+^ strains over a period of seven days. Within 24 hours, the viability of the WT strain was at 85%, whereas it was reduced to 70% for the *dksA^-^*strain and this trend continued for the entire course of the experiment (**Fig. 5A**). At the end of the study only 50% of the original population of *dksA^-^* strain survived, significantly less than the WT and the complement strain (**Fig. 5A**), implying that DksA influences *K. pneumoniae* environmental survival. To further elucidate the regulatory mechanisms underlying DksA-mediated environmental survival, we examined whether DksA affected the expression of the general stress response regulator RpoS, which was previously observed to contribute to *K. pneumoniae* survival on a solid surface (40) and other environmental stressors (55, 56). Furthermore, in *E. coli* DksA is known to regulate *RpoS* at the translational level(57, 58). Compared to the WT, we observed reduced RpoS levels in the *dksA^-^* strain (**Fig. S6A**), suggesting that DksA positive regulation of RpoS likely contributes to *K. pneumoniae* survival in the environment.

**Figure 5:**
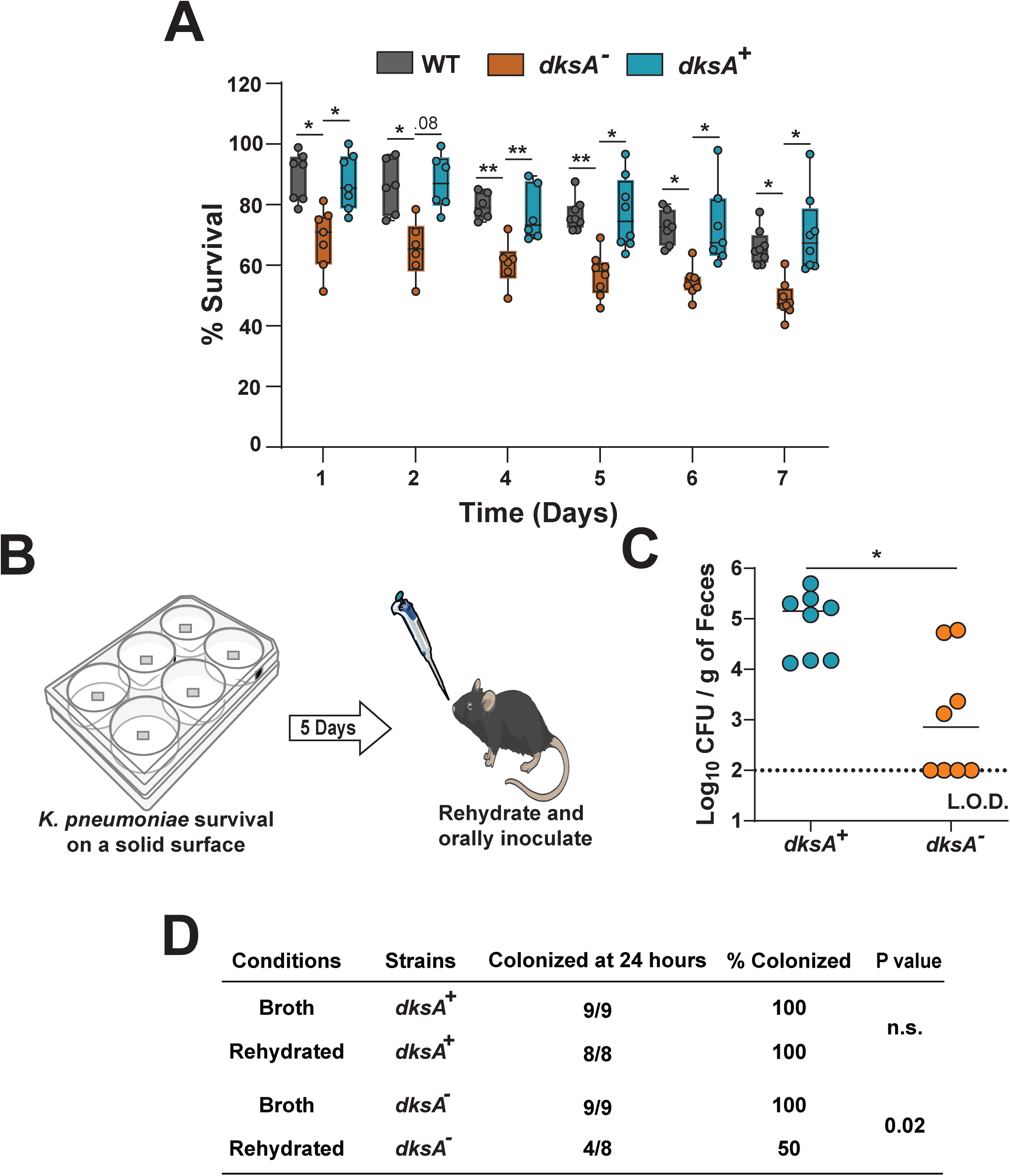
*K. pneumoniae* survival on a surface and subsequent acquisition by a naive host requires DksA. **(A)** *Ex vivo* solid surface survival of WT, *dksA*^-^, and *dksA^+^* strain. Bacterial strains were grown overnight, resuspended in PBS, OD_600_ adjusted to ∼4 and spotted onto nitrocellulose discs on 1% agarose pads in 6-well polystyrene plates. Discs were removed at the designated time points, washed with PBS, serially diluted and plated for viable bacterial counts. Data is presented as box-and-whisker with median and min to max shown for eight independent essays (in duplicate). Statistical analysis was carried out using Kruskal-Wallis tests with Dunn’s test of multiple comparisons at each time point. **(B)** Schematic representation of the protocol for *K. pneumoniae* acquisition after surviving on a solid surface and reinfection in mice. **(C)** Fecal shedding from mice after one day of infection with solid surface survived *K. pneumoniae* strains. Mice were orally inoculated (10^6^ CFU) either with *dksA*^-^ mutant, or the complement *dksA^+^*strain (*n* = 8 for each group) from bacteria residing on solid surface (nitrocellulose discs) for 5 days. Feces were collected 1-day postinfection to enumerate bacteria. Each point indicates a single mouse on a given day, the bars indicate the median shedding, and the dotted line indicates the limit of detection. The proportion of colonized mice were compared using Fisher’s exact test. (D) Comparison of acquisition of *K. pneumoniae* infection by naïve mice after 24 hours. Statistical significance was determined using Fisher’s exact test. *\*P* < 0.05; \**\**, *P <* 0.01; \**\*\**, *P <* 0.001; n.s., not significant.

Next, we investigated the impact of environmental survival on subsequent acquisition by a naïve host. Acquisition is an early step before establishment of colonization and is considered an important step in host-to-host transmission. *K. pnuemoniae* strains that had survived in the environment for 5 days were rehydrated and the CFU adjusted in PBS to 10^6^ CFU/100µl for all strains. We chose 24 h post inoculation as an arbitrary timepoint to determine successful acquisition of *K. pneumoniae*. Naïve mice were orally inoculated with the rehydrated samples and fecal shedding used to determine successful acquisition. We observed that all of the mice inoculated with *dksA^+^* from both the rehydrated (**Fig. 5B**) and LB grown (**Fig. 4A**) had successful acquisition (100%) at 24 h. However, compared to LB grown, the rehydrated *dksA^-^*strain had lower acquisition events (50%) (**Fig. 5C-D**). Compared to *dksA^+^* strain, the bacterial burdens in tissues from lower intestine were also significantly reduced in *dksA^-^* strains inoculated mice (**Fig. S6B**). Thus, our results suggest that in addition to desiccation tolerance DksA also contributes to acquisition of *K. pneumoniae* by a new host.

## Discussion

*K. pneumoniae* employs diverse regulatory networks to integrate metabolic flexibility with stress tolerance for persistence across both environmental reservoirs and mammalian hosts (59). DksA is an ancestral potentiator of the stringent response that modulates RNA polymerase activity to fine-tune transcriptional programs in response to environmental cues (13, 19). Our findings reveal a distinct functional role of DksA in *K. pneumoniae*, where it governs traits including growth under nutrient limitation, antibiotic resistance, mucoviscosity, biofilm formation, and survival on nutrient-depleted surfaces that are critical for persistence on abiotic surfaces. Notably, biofilm formation and desiccation tolerance have been shown to enhance long-term environmental survival of *K. pneumoniae* and preserve infectivity following environmental exposure (40, 53). Similar links between global stress regulators, environmental persistence, and colonization fitness have been described in other pathogens, underscoring the broader relevance of this regulatory strategy (25, 28, 29, 32, 60).

Our growth studies established that DksA promotes metabolic adaptation under nutrient limitation, as *dksA* deletion resulted in a growth defect in minimal medium that was alleviated by casamino acid supplementation (**Fig. S1B**, **C**). Elevated *dksA* promoter activity during exponential phase suggests that DksA acts early to adjust with nutrient limitation and likely promotes expression of genes involved in amino acid biosynthesis during nutrient scarcity observed in other Gram-negative pathogens (25, 27, 36). Beyond metabolic adaptation, The *dksA*^-^ strain exhibited increased resistance to aminoglycosides, tobramycin and kanamycin (**Fig. 1 B, C**), which is in line with results obtained with *V. cholerae* (38). In contrast, *dksA* deletion in *A. baumannii* resulted in increased sensitivity to aminoglycosides and no difference with cationic antimicrobial peptides (61). These contradictory results suggest that while DksA is conserved across Gammaproteobacteria, how it modulates gene expression during stringent stress response is nuanced. In addition, *dksA* deletion altered outer membrane permeability without detectable disruption of inner membrane integrity (**Fig. 1 E, F**). These observations suggest that loss of DksA alters outer membrane architecture or composition, potentially affecting lipid organization or surface charge, which could explain the increase in resistance observed with polymyxin B (**Fig. 1A**) whose primary target is the lipid A moiety of LPS. Additionally, aminoglycoside are also known to interact with LPS and porins present on the outer membrane (39, 62–64). Similar regulatory perturbations have been observed to reshape membrane structure and antibiotic susceptibility in other pathogens (34, 38). Recent molecular epidemiological data suggest that certain *K. pneumoniae* carbapenem resistant isolates also harbor deletion in *dksA*, indicating that *dksA* mutations can arise in a clinical setting, potentially because of selective antibiotic pressure (65). Then again, while *dksA* deletion confers antibiotic resistance it likely comes with a biological cost as demonstrated by our *in vitro* and *in vivo* studies.

Research work identifying regulatory factors that are conserved across both classical and hypervirulent *K. pneumoniae* strains and other Enterobacteriaceae has highlighted the importance of global transcriptional regulators such as H-NS, Fur, CpxR, Fnr and Hfq in shaping virulence-associated phenotypes independent of strain background (66–70). Capsule production and hypermucoviscosity (HMV) are often linked as defining virulence-associated traits of *K. pneumoniae* and several transcriptional regulators have been identified as contributing to capsule gene expression in *K. pneumoniae* strains (66, 71, 72). We established that *dksA* deletion effected the expression of two out of three *cps* locus promoters without affecting total capsule abundance (**Fig. 2 B-E**) suggesting that the genes downstream of the *wzi* promoter are likely the major drivers of *K. pneumoniae* capsule levels. These results differ from those obtained with *A. baumannii* where in comparison to the WT, a *dksA* deletion mutant had lower capsule levels (32). Subsequently, we focused on HMV, a capsule related but distinct phenotype, where the small protein RmpD interacts with capsular polysaccharide biosynthesis and export enzyme Wzc to produce more uniform CPS chain lengths (45, 73–75). We found that deletion of *dksA* reduced the expression of *rmp* locus promoter with a concurrent reduction in hypermucoviscosity (**Fig. 2 F-H**). Thus, our results add to the regulatory complexity of this critical *K. pneumoniae* pathogenesis determinant where *rmpA* positively regulates its own synthesis and that of *rmpD* and *rmpC.* In addition, ArgR, KvrA, KvrB, RcsB, and as illustrated, DksA, positively modulate the *rmp* locus and in turn HMV (72, 76), whereas Fur negatively regulates the *rmp* locus (77).

Besides CPS, biofilm formation is another well-established *K. pneumoniae* virulence phenotype, primarily associated with the type 3 fimbriae. DksA has been previously implicated in biofilm formation in *S. enterica* serovar Typhimurium (*S. Typhimurium*) and in *A. baumannii* with contrasting results. In *S. Typhimurium* DksA mediated positive regulation of *Salmonella* pathogenicity island 1 (SPI-1) genes that likely regulate biofilm formation, whereas in *A. baumannii* reduced CPS potentially contributes to the increase in biofilm formation (25, 78). Our static and shear flow biofilm studies provide a comprehensive understanding of the spatial organization of the biofilm components involved (**Fig. 3**) and our observations support a mechanistic framework in which DksA-mediated stringent response signaling likely integrates quorum sensing, and fimbrial expression to coordinate surface machinery and biofilm architecture of *K. pneumoniae*. Interestingly, while we observed reduced HMV, which generally negatively correlates with biofilm formation (46), we did not observe a corresponding increase in biofilm formation possibly because the reduction in expression of fimbriae and quorum sensing genes is much greater to overcome than the positive impact of HMV reduction.

Fomites are considered as a major source of continued *K. pneumoniae* fecal-oral transmission in a hospital setting (79, 80). Here, using our murine model of *K. pneumoniae* gut colonization, we establish that similarly to *S. Typhimurium* (25), *K. pneumoniae* DksA is required for gut colonization. Notably, this colonization defect persisted even after antibiotic-mediated disruption of the native gut microbiota (**Fig. 4D**), further dictating that DksA is required for colonization regardless of microbiota context. The inability of the *dksA^-^* strain to establish robust gut colonization likely reflects its compromised regulation of surface-associated and stress-adaptive traits that are critical for persistence within the gastrointestinal environment (7). Due to this intrinsic defect in gut colonization, we could not perform transmission studies with our *dskA^-^*strain using our murine model (52). However, as we observed poor survivability of the *dksA^-^*strain on a solid surface (**Fig. 5A**), mediated through RpoS, we postulated that DksA might contribute to acquisition, a key step of transmission (81). Our acquisition studies confirmed that DksA promotes *K. pneumoniae* acquisition (**Fig. 5C-D**). Thus, DksA as a master stress regulator influences different facets of the lifecycle of *K pneumoniae*, including gut colonization, survival outside the host and acquisition. In *A. baumannii* hydrophilins contribute to desiccation survival and in turn acquisition (82). Similarly, DksA besides RpoS, likely regulates other factors that contribute to survival outside the host. Future studies will aim to elucidate the genetic determinants that contribute to *K. pneumoniae* survival followed by acquisition to a new host.

In summary, our study experimentally advances our understanding of the role stringent response regulator DksA plays in modulating *K. pneumoniae* antibiotic resistance, membrane integrity, virulence-associated traits and gut colonization. In addition, DksA enables *K. pneumoniae* to survive in nutrient-limited environments and subsequent acquisition by a naïve host.

Environmental survival mechanisms used by pathogens and their impact on gut colonization remain understudied. These findings have a clinical relevance as *K. pneumoniae* MDR isolates are notoriously difficult to eradicate once they gain access to the hospital setting (79). Given the importance of environmental reservoirs in the transmission of nosocomial pathogens, targeting DksA-regulated pathways may represent a strategy to disrupt both environmental persistence and subsequent infection.

## Materials and Methods

### Ethics Statement

This study was conducted according to the guidelines outlined by the National Science Foundation animal welfare requirements and the Public Health Service Policy on Humane Care and Use of Laboratory Animals (83). All animal work was done according to the guidelines provided by the American Association for Laboratory Animal Science (AALAS) and with the approval of the Wake Forest Baptist Medical Center Institutional Animal Care and Use Committee (IACUC). The approved protocol number for this project was A20-084. Animal studies performed at Emory University School of Medicine were performed with the approval of IPROTO202500000047 by IACUC at Emory University.

### Bacterial strains and growth conditions

Bacterial strains, plasmids and primers used in this study are listed in **Table. 1** and **S1** respectively. All strains were grown aerobically in lysogeny both (LB; LB-Lennox) medium at 37°C unless otherwise stated. Where appropriate, antibiotics were added at the following concentrations: kanamycin (Kan) (50 µg/ml), streptomycin (Strep) (500 µg/ml), spectinomycin (Spec) (50 µg/ml), chloramphenicol (Cm) (50 μg/ml), ampicillin (Amp) (25 μg/ml) and apramycin (50 μg/ml).

### Strain and plasmid construction

To construct the *dksA* deletion mutant (*dksA*::*cam* [*dksA^-^*]), genomic DNA was isolated from the corresponding mutant with a transposon insert in *dksA* in the MKP103 background (84).

Approximately 500-bp homology on either end of the transposon element with the chloramphenicol (Cam) cassette was amplified using the Q5 polymerase (NEB). Purified PCR product was electroporated (1.8 kV, 400 Ω, 25 μF) into a KPPR1S-derivative containing the temperature-sensitive plasmid pKD46, containing the λ red recombination genes downstream of an arabinose-inducible promoter. Recombination was performed as described previously (85).

Successful mutants in the KPPR1S background were selected on LB agar with cam (50 μg/mL) and confirmed via colony PCR. Chloramphenicol cassette from *dksA::cam* was removed using the plasmid pCre2 as described previously to yield *ΔdksA* strain (86).

The *Δdks*A chromosomal complement (*dksA^+^*) was constructed as described previously (87). Briefly, the *dksA* gene and ∼500 bp upstream and downstream was amplified using KPPR1S genomic DNA via Q5 polymerase (NEB). Plasmid pKAS46 (**Table. 1**) was digested with NotI-HF and NheI-HF, and the purified PCR product was ligated using the NEBuilder HiFi DNA assembly kit (NEB; E2621L) and transformed into *Escherichia coli* S17-1 *λpir*. Successful transformants were confirmed via PCR. Conjugation with *ΔdksA* (**Table. 1**) strain and successful complemented strain was selected and confirmed as described previously (40).

To construct reporter fusions plasmids pPROBE-tagless-*gfp* plasmid (88) was used to insert the promoter region of *dksA* and *mrkA* gene upstream of *gfp* (transcriptional fusion) region. DNA fragment containing (∼500 bp) promoter region were amplified using KPPR1S genomic DNA via Q5 polymerase (NEB). The pPROBE plasmid was digested with SalI-HF and BamHI-HF restriction enzyme and assembled with the amplified *dksA* promoter region using the NEBuilder HiFi DNA Assembly Master Mix (NEB; E2621L). The assembled product was transformed into *Escherichia coli* S17-1 *λpir* and the resultant transformants were confirmed via PCR and Sanger sequencing. Subsequently, the plasmid was transformed into WT, *dksA^-^* and *dksA^+^* to examine the GFP levels *in vitro.* Primers used for making the *gfp* constructs are listed in **Table S1**.

To construct the constitutive *gfp* expressing strain, *Pem7-gfp* cassette region was amplified from pJH026 plasmid (89) and ligated with NsiI-HF (NEB), XhoI (NEB) digested pUC18R6K-mini-Tn7T-Km plasmid using NEBuilder master mix (NEB). The ligated product was then transformed into *E. coli* S17-1 *λpir*, selected on agar plates containing kan and positive colonies were confirmed by PCR and sequencing. The resulting plasmid was integrated at the *attTn7* site of *K. pneumoniae* strains with the help of pTNS2 plasmid by electroporation as described previously (90). Kanamycin resistant colonies were confirmed via PCR to determine the insertion and orientation of the fragment. *gfp* expression was further confirmed by Varioskan LUX Multimode Microplate Reader (Thermo Scientific).

### Growth curve

Bacterial growth was measured in three different media including LB, M9 Minimal Media (MM) with 0.4% Glucose (Glu; sole carbon source), and M9 MM with 0.2% casamino acids (CAA). Briefly, single colony from experimental strains (KPPR1S WT, *dksA^-^* and *dksA^+^*) was added to 5 mL LB and grown overnight at 37°C with constant agitation. Next day 1 ml culture from each tested strain was washed twice in 1X PBS, diluted 1:100 into 5 mL respective test media and grown at 37°C with constant agitation. At specific time points, a 20 μL sample was taken from each tube, serially diluted in 96-well plate and samples plated on selective antibiotic plates for colony count. Each strain was cultured on three independent days with three technical replicates.

### Antimicrobial resistance assay

Antibiotic resistance assays for polymyxin B (P4932-1MU, Sigma), tobramycin (455430050, Fisher), kanamycin (K1377, Sigma), cefixime (18588F, Sigma), nalidixic acid (N8878, Sigma), and oxacillin (AAJ67006AD, Fisher) were performed in 96-well microtiter plates as described previously (40). Briefly, overnight cultures of *K. pneumoniae* strains to be tested were grown in cation-adjusted Mueller-Hinton Broth (caMHB) broth then diluted 1:100 in fresh media and grown to OD_600_ of 1.0. Subsequently, cultures were diluted to ∼10^6^ CFU/mL in caMHB. 96-well plates were prepared with 100 μL of appropriate concentrations of tested antibiotics and 100 μL of the bacterial suspension was added to each well in triplicate. The plates were incubated at 37°C for 30 minutes (for cefixime the incubation time was 3 hours), 20 μL was removed from each well, serially diluted, and samples plated on selective antibiotic plates for enumeration.

### Membrane permeability assay

Membrane permeability analysis was performed as described previously (38) with slight modifications. Briefly, Bacterial strains were cultured overnight in LB and then diluted 100-fold in fresh LB and grown till they reached OD_600_ = 1.0. Subsequently, 1 ml culture was pelleted down (21,000 × g) at 4°C, washed twice with chilled 1X PBS, and resuspended in the same volume of PBS. Afterwards, in a flat bottom 96-well plate (Fisher, FB012931) 200 µL of bacterial sample was added in triplicates. For outer membrane permeability, 1-N-phenylnaphthylamine (NPN) was added to the bacterial samples at a final concentration of 20 µM. Fluorescent reading was taken immediately using Varioskan LUX Multimode Microplate Reader (Thermo Scientific) at an excitation of 350 nm and 420 nm emission respectively. For inner membrane permeability, samples were prepared as described above and propidium iodide (PI) used at a final concentration of 2.5 µg/mL. The reading was taken at an excitation wavelength of 535 nm and an emission of 617 nm. Assays were performed in triplicates with 3 biological replicates.

### Uronic acid measurement

Capsular polysaccharide amount was determined by quantifying the uronic acid content as described previously (44). Briefly, bacterial strains were grown overnight at 37°C with constant agitation, subcultured to an OD_600_ of 0.2. After 6 hours of incubation at 37°C, 500 µl of culture was mixed with 100 µl 1% zwittergent-100 mM citric acid and incubated at 50°C for 20 min.

The samples were centrifuged for 5 min at 20,000 x *g*, and 300 µl of supernatant was removed and precipitated with 4 volumes of absolute ethanol. The precipitated sample was centrifuged as above to pellet the precipitated material, the pellets were air dried and resuspended in 200 µl distilled water. 1.2 ml sodium tetraborate–concentrated H_2_SO_4_ were added in each sample, vortex vigorously and the reaction mixture boiled for 10 mins and placed on ice for 10 mins. Afterwards, 20 µl of 0.15% 3-phenylphenol (Sigma-Aldrich) that acts as a colorimetric indicator was added to the sample, incubated at room temperature for 5 min and the absorbance measured at 520 nm. A standard curve was used to detect glucuronic acid content using glucuronate lactone (Sigma-Aldrich, St. Louis, MO) and expressed as micrograms per 10^9^ cells.

### Mucoviscosity assay

The mucoviscosity assay was conducted using the sedimentation assay as previously described (91) with slight modifications. Briefly, Bacterial cultures were grown overnight in LB at 37°C and were subcultured to an OD_600_ of 0.2 in fresh LB media. After 6 hours, the cultures were set to an OD of 1.0 and centrifuged at 1,000 x *g* for 5 min. The supernatant was carefully removed and OD_600_ recorded. Final readings were normalized to the OD_600_ before centrifugation.

### Measurement of promoter activity

For time course assays, the test strains were grown overnight at 37°C with constant agitation in LB supplemented with 50 μg/mL kan. Overnight cultures were spun down and resuspend in fresh LB and adjusted to an OD_600_ =3 and subsequently diluted 1:100 in same media. 100 μL of each sample was loaded with 3 technical replicates into a 96-well flat bottom tissue culture plate (Fisherbrand; FB012931). 100 μL of fresh LB was added with 3 technical replicates as a control. The lid of the 96-well plate was treated with 0.2% Triton in 20% ethanol to prevent fogging. All strains were assayed in triplicate, and each assay was performed at least 3 times. The optical density OD_600_ and the fluorescence of GFP at OD_540_ (RFU) of cultures at designated time points were measured using Synergy H1 Microplate Reader (BioTek). RFU values were normalized to OD_600_. To determine promoter activity of the *cps* and the *rmp* locus, strains to be tested were grown overnight as described above and subcultured to an OD_600_ of 0.2 in fresh LB media. After 6 hours of growth at 37°C, the optical density OD_600_ and the fluorescence of GFP at OD_540_ (RFU) of cultures were measured and the RFU values were normalized to OD_600_.

### Crystal Violet Biofilm Staining

Overnight cultures were grown in LB for 24 hours in shaking incubator (220 rpm) at 37°C. Overnight cultures were diluted to an OD_600_ of 0.5 (9.75 × 10^9^) in biofilm media (tryptic soy broth and 0.5% glucose). Two mL of standardized culture was seeded into tissue culture treated 6-well plates and incubated statically for 24 hours at 37°C. The supernatant was removed and samples were washed with 1 mL of 1× PBS. Biofilms were stained by adding 2 mL 0.1% crystal violet for 15 minutes. 0.1% crystal violet was removed, and 6-well plates were dried in fume hood for 24 hours. Biofilms were de-stained by adding 2 mL 30% acetic for 15 minutes, followed by transferring samples to a 96-well plate. SYNERGY H1 Microplate reader (BioTek) was used to measure the optical density at 550 nm. Control wells were seeded with sterile biofilm media, incubated for 24 hours, and stained alongside the biofilm samples. The OD_550_ values obtained from the control wells were subtracted from OD_550_ values of biofilm samples to normalize and account for dye binding to the plastic wells. All strains were tested in biological and technical triplicates.

### Static Growth Confocal Microscopy Biofilm Imaging

As described previously (46), cultures of chromosomally encoding *gfp K. pneumoniae* strains were grown and standardized as above. 1 mL of the standardized culture was pipetted into Matsunami glass dishes, sealed with Parafilm, and placed in 37°C static incubator for 24 hours. After 24 hours, supernatant was removed, and samples were washed with 1 mL of 1× PBS. To assess the presence of lipids, 50 µg mL^-1^ Nile red dye in molecular-grade water was added and the dish transferred to a rocker on medium speed for 40 minutes followed by removal of the stain and washing of samples with 1 mL of 1× PBS. To visualize biofilm matrix polysaccharides, 50 µg mL^-1^ calcofluor white dye in molecular-grade water was pipetted onto samples and incubated for 5 minutes on a rocker. Calcofluor white stain was removed, and samples were washed with 1 mL of 1× PBS. 3D-rendered z-stack confocal microscopy was used to image biofilm samples.

### Shear Flow Confocal Microscopy Biofilm Imaging

To assess biofilm formation capabilities under shear flow conditions, the BioFlux One system (Cell Microsystems, Durham, NC) was utilized. Cultures were grown in biofilm media as described above. The BioFlux One system was powered on and heated to 37°C along with the biofilm media and 48-well low shear plate (0–20 dyne/cm²). 40 μL of warm media was pipetted into the outlet wells and perfused from outlet to inlet wells at 20 dyne/cm² (1874 μL h^−1^) for 40 seconds. To avoid drying of the inlet wells, 100 μL of warm media was added and residual media was removed from the outlet wells. 20 μL of standardized culture was added to the outlet wells and perfused from outlet to inlet at 5 dyne/cm² (468 μL h^−1^) for 5 seconds. The 48-well low shear plate (0–20 dyne/cm²) was incubated for 1 hour at 37°C without flow. 1 mL of warm media was used to rinse the outlet wells and 1 mL warm media was pipetted into the inlet wells and perfused at 1 dyne/cm² (94 μL h^−1^) for 5 minutes from inlet to outlet wells. Outlet wells were rinsed with 1 mL warm media and inlet wells were assessed to ensure 1 mL media was still present. 48-well low shear plate (0–20 dyne/cm²) received flow from inlet to outlet wells at 0.69 dyne/cm² (65 μL h^−1^) for 24 hours. After 24-hour shear flow, wells were cleared of liquid and 100 μL of bacterial suspension in 1× PBS was pipetted into the inlet wells and perfused at 0.99 dyne/cm² (93 μL h^−1^) for 15 minutes to visualize cells. Wells were cleared of liquid and 100 μL of 1× PBS was added to the inlet wells and perfused from inlet to outlet wells at 0.99 dyne/cm² (93 μL h^−1^) for 15 minutes. To assess lipids, 50 µg mL^-1^ Nile red dye in molecular-grade water was added to inlet wells and perfused from inlet to outlet wells at 0.99 dyne/cm² (93 μL h^−1^) for 15 minutes followed by clearing of the wells and flowing of 100 μL of 1× PBS for 15 minutes at same flow rate. Biofilm polysaccharides were stained with 50 µg mL^-1^ calcofluor white prepared in molecular-grade water and 100 μL was added to inlet wells and flowed from inlet to outlet wells at 0.99 dyne/cm² (93 μL h^−1^) for 15 minutes. 100 μL of 1× PBS was flowed from inlet to outlet wells at same flow rate to wash channels. BioFlux One 48-well low shear plate (0–20 dyne/cm²) was imaged using z-stack confocal microscopy.

### Confocal Microscopy Imaging

Biofilm samples were imaged using z-stack confocal microscopy imaging with a Zeiss LSM 710 confocal microscope and a 63 × oil immersion lens. Lens oil was applied to the microscope lens and sample dishes were placed on the microscope lens holder. 488 nm laser channel was used to image calcofluor white while 543 nm laser was used for GFP and Nile red stains. Images were captured using confocal z-stacks (0–30 µm) with 3D renderings of the z-stack images generated using ZEN microscope software.

### RNA extraction, cDNA synthesis, and qRT-PCR

RNA was isolated from *in vitro* gown samples using TRIzol reagent as described previously (44). Briefly, Bacterial cultures were grown overnight in LB and then subcultured (1:100) in fresh LB and grown to OD_600_ =1.0 followed by RNA isolation. Total RNA was treated with DNase (Invitrogen; AM1907) followed by cDNA synthesis using an iScript cDNA synthesis kit (iScript™, BIO-RAD). qRT-PCR was conducted with 20 ng cDNA and 0.5 mM primers per reaction using iTaq Universal SYBR green Supermix. (Bio-Rad; 1725121) *K. pneumoniae* species-specific primers listed in **Table. S1** was used and *gyrA* was used as internal control. Expression threshold cycle (*C_T_*) was quantified via the (2*^−ΔΔC^_T_*) method (92). The data were shown from an average of 3 biological replicates that were grown and processed for RNA isolation on separate days.

### Mouse infections for shedding and colonization

All mice infections experiments were conducted as previously described (52, 93). Briefly, specific-pathogen-free (SPF) C57BL/6J mice were purchased from Jackson Laboratory (Bar Harbor, ME) and bred and maintained in the animal facility at Biotech Place at Wake Forest School of Medicine. 6–8-week-old mice had their food and water removed for 3 hours before bacterial inoculation. Mice were orally fed an infectious dose of ∼10^6^ CFU/100 μl of *K*. *pneumoniae* diluted in PBS with 2% sucrose in two 50-μL doses an hour apart. To enumerate the infection dose the inoculation samples were serially diluted and plated for CFU count.

GI colonization was determined by fecal collection and quantification as described previously (52). Briefly, fecal pellets (approximately 2 pellets) were collected, weighed, diluted 1:10 in PBS (weight/volume) and homogenized with at least 2 glass beads using a bead mill homogenizer (Fisherbrand; 15-340-163). Samples were then briefly spun using a microcentrifuge (Thermo Scientific; MySpin 6) and the supernatant was serial diluted in PBS and plated on selective antibiotic plates for enumeration. At the experimental endpoint, mice were euthanized with CO_2_ exposure (2 liters/minute for 5 minutes) followed by cardiac puncture. To determine bacterial density in the oropharynx and the lower GI tract, oropharyngeal lavage and ileum, cecum, and colon tissue were harvested, homogenized, serially diluted and plated on selective antibiotic plates for enumeration (limit of detection: 10^2^ CFU/mL).

For competition studies (CI), an apramycin resistant derivative of KPPR1, *dksA^-^*, *dksA^+^* were used to prepare infection dose containing a 1:1 ratio. Mice were fed with inoculation dose of ∼1×10^6^/100 μL as described above. Fecal pellets and organs were collected as described above and homogenates, diluted and plated on both apramycin (50 μg/ml) and strep (500 μg/ml) LB agar plates to enumerate the colonization. The CI was calculated by using the following equation:

log^10^ CI = (mutant output over WT output) / (mutant input over WT input)

To induce antibiotic dependent supershedder phenotype mice were infected with KPPPR1S as described above and subsequently orally gavaged with a single dose of strep (5 mg/200 μL in sterile PBS) 2 days post inoculation. Daily shedding was monitored post-antibiotic treatment for an additional 5 days.

Surviving bacteria were harvested from nitrocellulose membranes and rehydrated and washed twice with 1X PBS and resuspended in PBS with 2% sucrose for a final concentration ∼10^6^ CFU/100 μl. Mice were orally inoculated as described above. Bacterial colonization was determined by fecal collection and quantification as described.

### Solid surface survival assay

Bacterial survivability on a solid surface in the absence of nutrition was determined as previously described (40). Briefly, strains to be tested were grown overnight in LB. The cultures were then centrifuged at 21,000 × *g* for 25 minutes, washed and resuspend in PBS to adjusted OD_600_ of 4.0. 1% agarose (Fisher; BP160-500) was added to a 6-well polystyrene plate (Costar; 3516) to create pads, and after solidifying nitrocellulose membranes (MF-Millipore; HAWP02500) were placed over the agarose pads. Afterwards, twenty microliters of bacterial suspensions were spotted on the membrane and air dried for 20 minutes, followed by covering the plate with a lid and parafilm to maintain moisture. For sampling, individual membrane was removed, washed in 1 mL PBS to dislodge the adhered bacteria, serially diluted and plated onto selective LB agar plates. In between sampling, plates were sealed with parafilm (Bemis; PM-999) and stored at room temperature in the dark.

### Western blotting

Western blot was performed as described previously with slight modifications (40, 94). Briefly, Bacterial strains were grown in caMHB overnight. A total of 500 μL overnight culture was pelleted and resuspended in 1X Laemmli sample buffer (Bio-Rad) using the formula OD_600_/6 × 250 and boiled for 20 minutes. Samples were separated in 4–20% Mini-PROTEAN® TGX™ Precast Protein Gels (Bio-Rad; 4561093) and transferred to a nitrocellulose membrane (Bio-Rad; 1704158) using the Bio-Rad Trans-Blot Turbo transfer system. The membrane was blocked in 1X Tris-Buffered Saline (Bio-Rad; 1706435) with Tween-20 (0.1%) (TBST) containing 5% nonfat dry milk (Bio-Rad;1706404) at room temperature for 1.5 h and incubated with primary antibody anti-RNA sigma S antibody (1:4000) (BioLegend; cat. number 663703) and anti-*E. coli* RNA Polymerase β Prime Antibody (1:4000) (BioLegend; cat. number 662904) overnight at 4°C. Subsequently, the membranes were washed with TBST three times and incubated with secondary antibody goat anti-mouse IgG (H+L)-horseradish peroxidase (HRP) conjugate (Bio-Rad; cat. number 1706516) (1:10000) at room temperature for 2 hours. After washing with TBST, the blot was developed with Clarity western ECL substrate (Bio-Rad; 170-5061). Immune reactive bands were visualized by Bio-Rad ChemiDoc MP imaging system, and blot analysis was conducted using ImageJ software. (95)

### Statistical analysis

All statistical analyses were performed using GraphPad Prism 10 (GraphPad Software, Inc., San Diego, CA). Comparisons between two groups were analyzed via the Mann-Whitney *U* test, comparisons between multiple groups were analyzed via the Kruskal-Wallis test with Dunn’s post-analysis, and competitive index/competition experiments were analyzed using the Wilcoxon signed-rank test. Fisher’s exact test was used to determine significance between two categorical variables.

## Acknowledgement

We thank Dr. Kimberly A Walker (UNC Chapel Hill) for the pPROBE plasmids described in this study, and for providing critical feedback on this manuscript. This study was funded by the Burroughs Wellcome Fund GDEP Award, NIAID training funds, and a diversity supplement award for G.E.H (T32AI007401; AI173244) and NIAID funding (AI166642; AI173244) to M.A.Z and R00AI163295 to R.M.F.

## Figure legends

**Figure S1: DksA is required for robust growth in minimal medium and positively regulates its own expression.**

**(A-C)** Growth kinetics of *K. pneumoniae* KPPR1S, the isogenic *dksA* mutant, and chromosomal complement *dksA^+^*in nutrient rich LB-Lennox media **(A)** or M9 minimal medium containing 0.4% glucose as a carbon source **(B)** or M9 minimal medium supplemented with 0.2% casamino acid (CAA) **(C)** at 37°C, under aerobic growth conditions. The growth kinetics were depicted by CFU/mL over time. Data presents the mean ± SEM of 3 independent experiments in duplicate. **(D)** GFP kinetic assay of the WT and *dksA^-^* mutant strain carrying plasmid with *dksA* promoter transcriptional *gfp* fusion (*dksA΄-gfp^+^*), with empty vector(pPROBE) as a reference. Data was obtained at 2-hour intervals for 12 hours and is shown as RFU/OD_600_.

**Figure S2: *K. pneumoniae* resistance against quinolone and penicillin is not DksA dependent**

**(A-B)** WT, *dksA*^-^, and *dksA^+^*strains were grown until OD_600_=∼1.0 in cation-adjusted MHB. Diluted samples (∼10^6^ CFU/mL) were mixed with increasing concentration of Nalidixic acid **(A),** Oxacillin **(B)** and incubated for 30 minutes at 37°C. Shown is the mean and SEM of three independent assays (in duplicate).

**Figure S3: *K. pneumoniae* biofilm formation, type 3 fimbriae (*mrk* locus), quorum sensing (*lsr* locus) genes are positively regulated by DksA.**

**(A)** Comparison of biofilm biomas formed on 6-well polystyrene plates when WT, *dksA^-^* and the complement strain *dksA*^+^ were grown statically in tryptic soy broth and 0.5% glucose at 37°C for 24 h by crystal violet staining assay (top panel) and quantification (bottom panel). Data is mean and ± SEM for three biological repeats. **(B)** Genetic organization of the type 3 fimbriae locus (top panel). The expression of genes highlighted in red and green was determined. Quantitative real-time PCR (qRT-PCR) comparing *mrk* gene expression between the WT and its isogenic *dksA* mutant when grown in LB-Lennox (OD_600_ = 1.0). Displayed is fold change in transcription of *mrkA, mrkB* genes between WT and mutant. gyrase (*gyrA*) was used as the housekeeping gene for calculating 2^−*ΔΔC*^*_T_*. Bars represent mean ± SEM for 3 biological replicates. Statistical differences were calculated using the Mann-Whitney *U* test. **(C)** *gfp* expression assay using WT, *dksA*^-^, and *dksA^+^* strains carrying plasmid with promoter region from *mrkA* promoter fused to *gfp* (pPROBE-*mrkA*′*-gfp^+^*). Data was obtained after 18h of incubation and the relative fluorescent units (RFU) were normalized to OD_600._ Data are represented from 3 independent assays. To determine statistical differences a Kruskal-Wallis followed by Dunn’s test of multiple comparisons was performed. **(D)** Genetic organization of quorum sensing genes *lsr* locus (top panel) with *lsrR* and *lsrA* highlighted in red and green as their expression was determined. WT and the isogenic *dksA^-^* mutant were grown in LB until OD_600_=1 and gene expression compared for *lsrR* and *lsrA* through qRT-PCR. Data is presented relative to WT strain with DNA gyrase (*gyrA*) serving as housekeeping gene for calculating 2^−*ΔΔC*^*_T_*. Bars represent mean ± SEM for 3 biological replicates. Statistical differences were calculated using the Mann-Whitney *U* test. *, *P* < 0.05; **, *P* < 0.01; ***, *P* < 0.001; n.s.; not significant.

**Figure S4: DksA regulates biofilm architectural components.**

Confocal z-stack imaging of constitutively *gfp* expressing *K. pneumoniae* WT (top panel), *dksA*^-^(middle panel), and *dksA^+^* (bottom panel) strains, Nile red was used to visualize the presence of lipids (red) and calcofluor white (blue) used to visualize the polysaccharide matrix, respectively. Each strain is shown grown in biofilm media under static conditions. For each strain the z-stack images are shown to visualize the height of the biofilm and images from the top of the biofilm are shown to visualize the overall cellular density. Biofilms were grown and imaged in triplicates with a representative image shown.

**Figure S5: DksA contributes to *K. pneumoniae* gut colonization.**

**(A)** Colonization density of the WT, *dksA^-^* and *dksA^+^* within Ileum, cecum, colon and oropharynx 7 days postinfection. Mice were euthanized, tissues homogenized and plated for bacterial enumeration. Each symbol represents single mouse. The dotted line indicates the limit of detection. Kruskal-Wallis test followed by Dunn’s test of multiple comparisons was performed for statistical analysis. **(B-C)** Fecal shedding on the indicated days **(B)**, and colonization of ileum, cecum, colon and oropharynx at day 7 postinfection **(C)**, from *in vivo* competitive index (CI) studies. Mice were orally infected with a 1:1 mixture of WT and *dksA^+^ K. pneumoniae* (n =10). Each point represents the log_10_ CI value from an individual mouse on the indicated day or at different organs and the dotted line indicates a CI of 1. One sample Wilcoxon signed rank tests were performed for each group against a theoretical value of 0. *, *P* < 0.05; **, *P* < 0.01; n.s.; not significant.

**Figure S6: Inactivation of *dksA* impacts *K. pneumoniae* survival outside of host.**

**(A)** Western blot analysis of RpoS levels in whole-cell extracts grown in cation-adjusted MHB until OD_600_=1. The RNA polymerase β′ subunit was used as a loading control. ImageJ was used to analyze the data. Shown is mean with ± SEM from four independent experiments. Kruskal-Wallis tests with Dunn’s test of multiple comparisons were used to determine statistical significance. **(B)** The mutant (*dksA*^-^) and the chromosomal complement strain (*dksA*^+^) were harvested from nitrocellulose discs 5 days after spotting and resuspended in PBS. The CFU was adjusted 10^6^ CFU/100µl for both strains, which was then used to orally inoculate mice. Colonization of *dksA^-^*and *dksA^+^* in the cecum, ileum and colon was determined at 7 days post inoculation and presented as percentage. Statistical significance was determined using Fisher’s Exact test. *, *P* < 0.05; **, *P* < 0.01.

---

**Table S1. Primers used in the study**

